# Discovery of Drought-Responsive Transposable Element-Related Peach miRNAs

**DOI:** 10.1101/143115

**Authors:** Asli Kurden Pekmezci, Gökhan Karakülah, Turgay Unver

**Affiliations:** İzmir International Biomedicine and Genome Institute (iBG-izmir), Dokuz Eylül University, 35340, İnciraltı, İzmir, Turkey; Present address: Egitim Mah., Ekrem Guer Sok. 26/3, 35340, Balcova, Izmir, Turkey

**Keywords:** microRNA, repetitive element, TE-miRNA, drought, *Prunus persica*

## Abstract

MicroRNAs (miRNA) are small non-coding regulatory RNAs that suppress their specific target transcripts either by cleavage or inhibition of translation. Transposable element related miRNAs (TE-miRNA) have been subjected to various studies so far, and some of them were found to be involved in stress response in plants. Here, small RNA (sRNA) sequencing libraries generated from drought stress treated peach tissues were utilized to identify TE-miRNAs. Our computational analysis led to the identification of 63 TE-miRNAs, which were either locating nearby of any TE (TE-related), or overlapping with a TE (TE-derived). Furthermore, 13 out of 63 TE-miRNAs were designated as drought-responsive. Expression pattern of the identified drought-responsive TE-miRNAs are observed as tissue-specific manner, and their specific target transcripts are mostly related to transcription factors and growth-associated genes. Our findings suggest that miRNAs in relation with transposable elements might be key molecular players in the regulation of drought response.

## INTRODUCTION

Transposable elements (TE) are DNA fragments that can move from one position to another in the host genomes, and they constitute the majority of plant genome (.Deragon et al. 2008). TEs are divided into two groups in plants, each group includes autonomous and non-autonomous elements (.Kejnovsky et al. 2012). Class I elements, retrotransposons, ‘copy and paste’ themselves by reverse transcription. The members of this group are LTR (Long Terminal Repeat), LINEs (Long Interspersed Elements) and SINEs (Short Interspersed Elements). LTR type retrotransposons include *copia* and *gypsy* groups, TRIM (Terminal-repeat Retrotransposons in Miniature) and LARD (Large Retrotransposon Derivatives) families (.Deragon, Casacuberta and Panaud 2008, Kejnovsky, Hawkins and Feschotte 2012). Class II elements ‘cut and paste’ themselves by excising from the chromosome and relocating at a new position. MITEs (Miniature inverted-repeat transposable elements) are the most common types of class II non-autonomous elements including TIR (Terminal inverted repeats) and Helitrons (.Lisch 2013, Wicker et al. 2007).

TEs play crucial roles in plant genome evolution by activating-deactivating or duplicating genes or creating entirely new ones because of their mutagenic potential. (.Bennetzen and Wang 2014). Movement of transposable elements can also leads to evolution of plant miRNA encoding genes (.Piriyapongsa and Jordan 2008). It was found that various number of miRNA genes were mobilized by transposable elements in grass genome (.Abrouk et al. 2012). The previous reports also demonstrated the association between miRNA gene evolution and TEs in *Arabidopsis thaliana* (.Maher et al. 2006, Piriyapongsa and Jordan 2008, Santiago et al. 2002).

MiRNAs are small (19-24 nt long), endogenously expressed non-coding RNAs which negatively regulate gene expression at post-transcriptional level (.Eldem et al. 2013). Besides their roles in plant development (.Mallory et al. 2005, Mallory et al. 2004, Palatnik et al. 2003), miRNAs play important roles in stress responses such as drought (.Qin et al. 2011). As one of the major abiotic factors, drought or water deprivation limits plant growth, crop yield, and quality (.Zhang 2015). The functional regulatory roles of miRNAs in drought stress response were reported in a number of studies conducted on model organisms such as barley (*Hordeum vulgare*) (.Kantar et al. 2010) wheat (*Triticum aestivum*) (.Akdogan et al. 2016), cotton (*Gossypium hirsitum*) (.Xie et al. 2015), switchgrass (*Panicum virgatum*) (.Xie et al. 2014), tobacco (*Nicotiana tabacum*) (.Frazier et al. 2011) and rice (*Oryza sativa*) (.Zhao et al. 2007). Functional studies with miR408 in chickpea (*Cicer arietinum*) (.Hajyzadeh et al. 2015), miR169 in tomato (*Solanum lycopersicum*) (.Zhang et al. 2011) and miR319 in creeping bent grass (*Agrostis stolonifera*) (.Zhou et al. 2013) proved the role of miRNAs in drought regulation TE-related miRNAs have been subjected to several studies so far. All the miRNAs located on repetitive elements in four plant species (*A. thaliana*, *Populus trichocarpa, O. sativa*, and *Sorghum bicolor*) were revealed in a previous report by Sun et al. (2012) (.Sun et al. 2012). In another study, TE-miRNAs in rice genome were discovered, and the question how the transposable elements are domesticated into new miRNA genes, was elucidated (.Li et al. 2011). Yongsheng, et al. (2011) identified two short-interfering RNAs (siRNA) as repetitive element derived and found out that these two small RNAs regulate abscisic acid signaling and abiotic stress response positively (.Yan et al. 2011). Therefore, TE-miRNA and drought stress response regulation in plants is one of the novel biological perspectives to be deeply analyzed for further translational applications.

Our group previously introduced drought-responsive set of miRNA sequences and their measurement at genome-wide in peach (*P. persica*) by high-throughput sequencing of control and drought tissues (.Eldem et al. 2012). In this study, we aimed to annotate TE-derived and TE-related peach miRNAs from the previously generated sRNA deep-sequencing libraries, and to identify drought-responsive ones among all TE-derived and TE-related miRNAs.

## MATERIALS AND METHOD

### miRNA identification

In this study, sRNA libraries generated in our previous study (.Eldem, Celikkol Akcay, Ozhuner, Bakir, Uranbey and Unver 2012) were utilized to identify TE-related miRNAs. Our sRNA dataset were consisting of four libraries generated from (i) roots of control sample (RC), (ii) roots of drought applied sample (RS), (iii) leaves of control sample (LC), and (iv) leaves of drought applied sample (LS). To identify and count the miRNAs in the libraries, *P. persica* genome (assembly: Prupe1_0) in multi-fasta format were downloaded from Ensembl Plants database v31 (http://plants.ensembl.org) (.Kersey et al. 2016). Furthermore, *P. persica* mature miRNA sequences was extracted as General Feature Format (GFF) file from miRBase release 21 (http://www.mirbase.org/) (.Kozomara and Griffiths-Jones 2014), and our sRNA libraries in fasta format were analyzed using miRDeep2 v2.0.0.7 (.Friedländer et al. 2012) with default parameters. The count values of the identified peach miRNAs were obtained as count per million (CPM) values. In order to detect differentially expressed miRNAs in control and stress libraries, we calculated fold change for each miRNA, and miRNAs with differential expression values ≥ 2 fold-changes we considered as drought responsive.

### Annotation of TE-derived and TE-related miRNAs

To determine genomic locations of the miRNAs identified by miRDeep2, we utilized BLAST tool v2.5.0. Known precursor-miRNA (pre-miRNA) sequences provided in miRDeep2 output were scanned within the peach genome using BLAST tool, and genomic location(s) of each pre-miRNA were determined. Afterwards, TE annotation, available at Ensembl Plants database, were downloaded, and examined whether TEs were co-located or overlapping with Bedtools v2.25.0 (.Quinlan 2014). We considered miRNA sequence as TE-related, if the distance between miRNA and TE was less than 1 kb. If miRNA sequence was overlapping >50% with any TE, it was classified as TE-derived.

### Target transcript prediction and gene ontology analysis

The target transcripts of TE-derived and TE-related miRNAs were predicted by an online miRNA target analysis tool; psRNATarget (http://plantgrn.noble.org/psRNATarget/) (.Dai and Zhao 2011). All TE-derived and TE-related miRNA sequences were uploaded to psRNAtarget web server, and their potential targets were extracted. In order to identify the functional annotation of the targets, Gene Ontology (GO) analysis of the target transcripts was performed with Blast2GO tool (.Conesa et al. 2005). The target transcript sequences were loaded to the tool in fasta format. BlastX search was done wit a parameter of e value as 1e-10. Then, GO terms of each Blast hit were obtained and these terms were categorized in three GO categories: (i) cellular component, (ii) molecular function, and (iii) biological process. The schematic representation of the methodology is shown in Figure 1.

**Figure 1.**
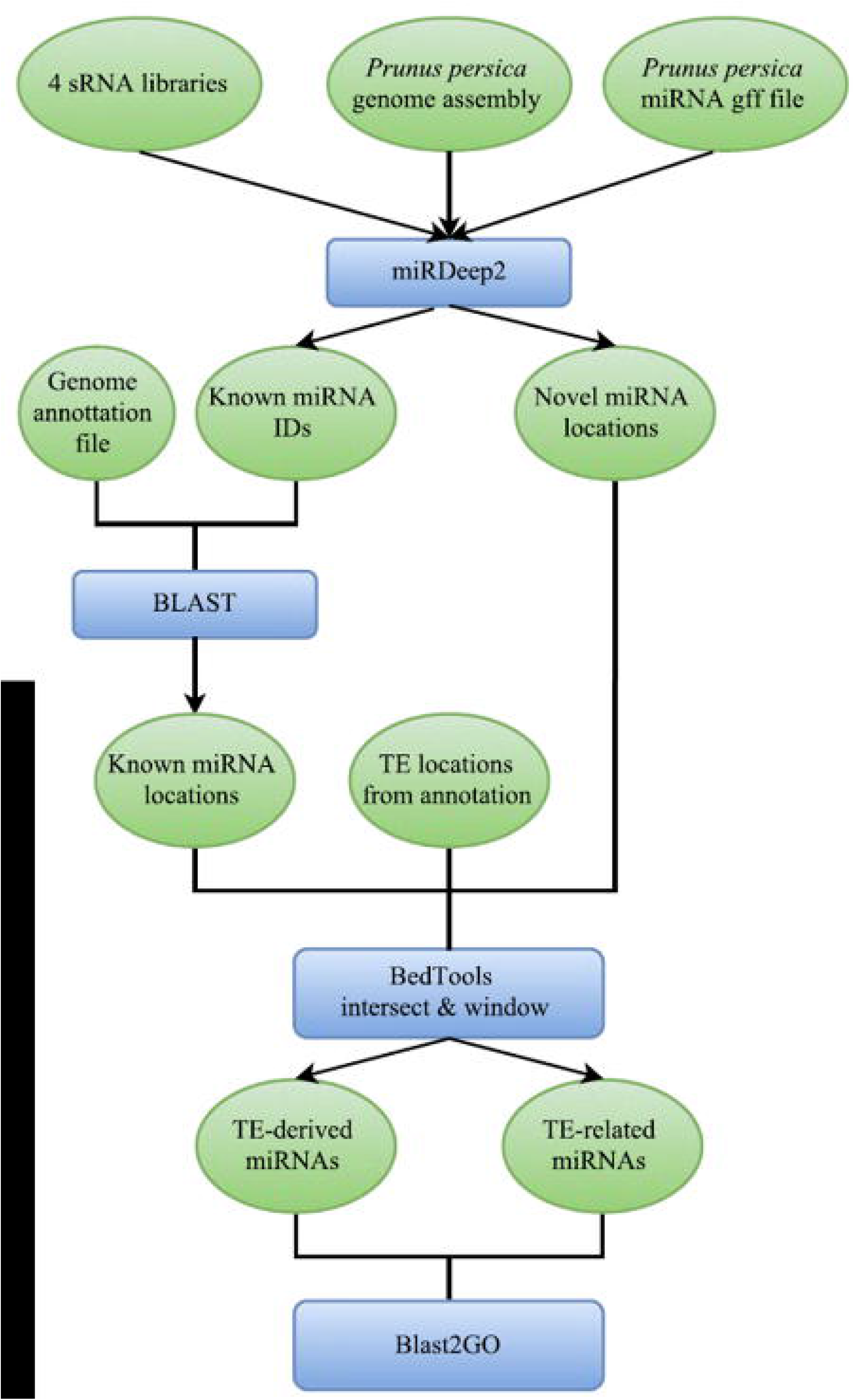
Flowchart representation of methodology. Blue rectangles represent tools, green ellipses are data used in tools as inputs.

## RESULTS

A total of 214 known miRNAs and 17 novel miRNAs were detected in *P. persica* genome by miRDeep2. Among the identified peach miRNAs that were found in at least one of the libraries, 63 miRNAs were defined as TE-miRNA (either TE-derived or TE-related) by the localization analysis. One of these miRNAs, ppe-MIR6284, was named as TE-derived miRNA. Seventy-two out of 97 nucleotide of this miRNA is in the repeat region named; LTR_Sl_chr_06_1092 according to Ensembl annotation (Figure 2).

**Figure 2.**
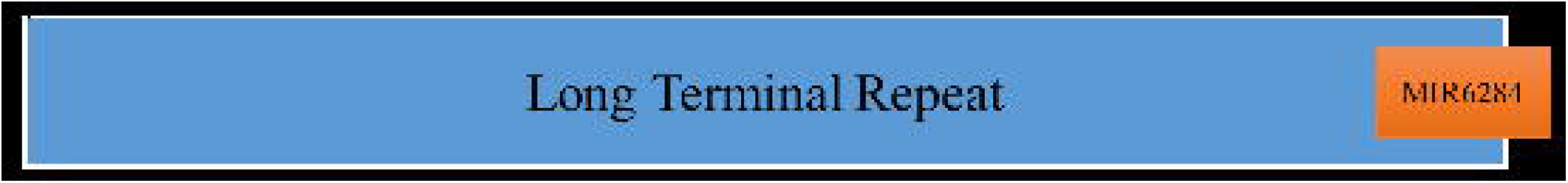
74 nucleotides of overlapping of MIR6284 with the LTR in chromosome 8, plus strand. Blue and orange blocks represent LTR_Sl_chr_06_1092 and ppe-MIR6284 respectively. The LTR begins at 20,947,395^th^ nucleotide and ends at 20,948,172^th^ nucleotides. ppe-MIR6284 gene is located in between 20,948,100^th^ and 20,948,196^th^ nucleotides of chromosome 8.

A total of 62 miRNAs was detected as TE-related and 58 of them were previously annotated known mRNAs such as miR166, miR171, miR172, miR399, and miR482. Besides, 4 of the 17 novel miRNAs were detected as TE-related (Supplementary Table 1). Most of TE-related miRNAs are in the window with more than one transposable element. We also identified 7 TE-related pre-miRNAs that were not detected in any of four sRNA sequencing libraries; MIR399a, MIR6267c, MIR6288c, MIR6291b, MIR6296, MIR7122a, and MIR828.

The comparative expression measurement analysis indicated that 58 of TE-related miRNAs were not differentially regulated in response to drought (Figure 3). Twelve of TE-related and one TE-derived miRNAs were detected as drought responsive with their differential expression pattern compared to stress treated versus control samples. Following TE-related miRNAs are reported as drought responsive; miR160b, miR162, miR167c, miR169g, miR169l, miR172a-5p, miR172d, miR5225-5p, miR6266a, miR6288c-3p, miR8123-5p, miR827 and miR6284 (Table 1). The expression values of miR6284, which is a drought responsive TE-derived miRNA, suppressed in leaves and up-regulated in roots with the fold changes of -2.52 (LC vs LS) and 2.36 (RC vs RS), respectively. The expression levels of most drought responsive TE-related miRNAs have changed in roots rather than leaves. Among them, the expression levels of miR169g and miR169l are seem to be changed in response to drought in roots, the former is related with an LTR, the latter is with a GA repeat. Fold change of miR162 is significant in roots, and miR6266a has differentially expressed both in leaves and roots.

**Figure 3.**
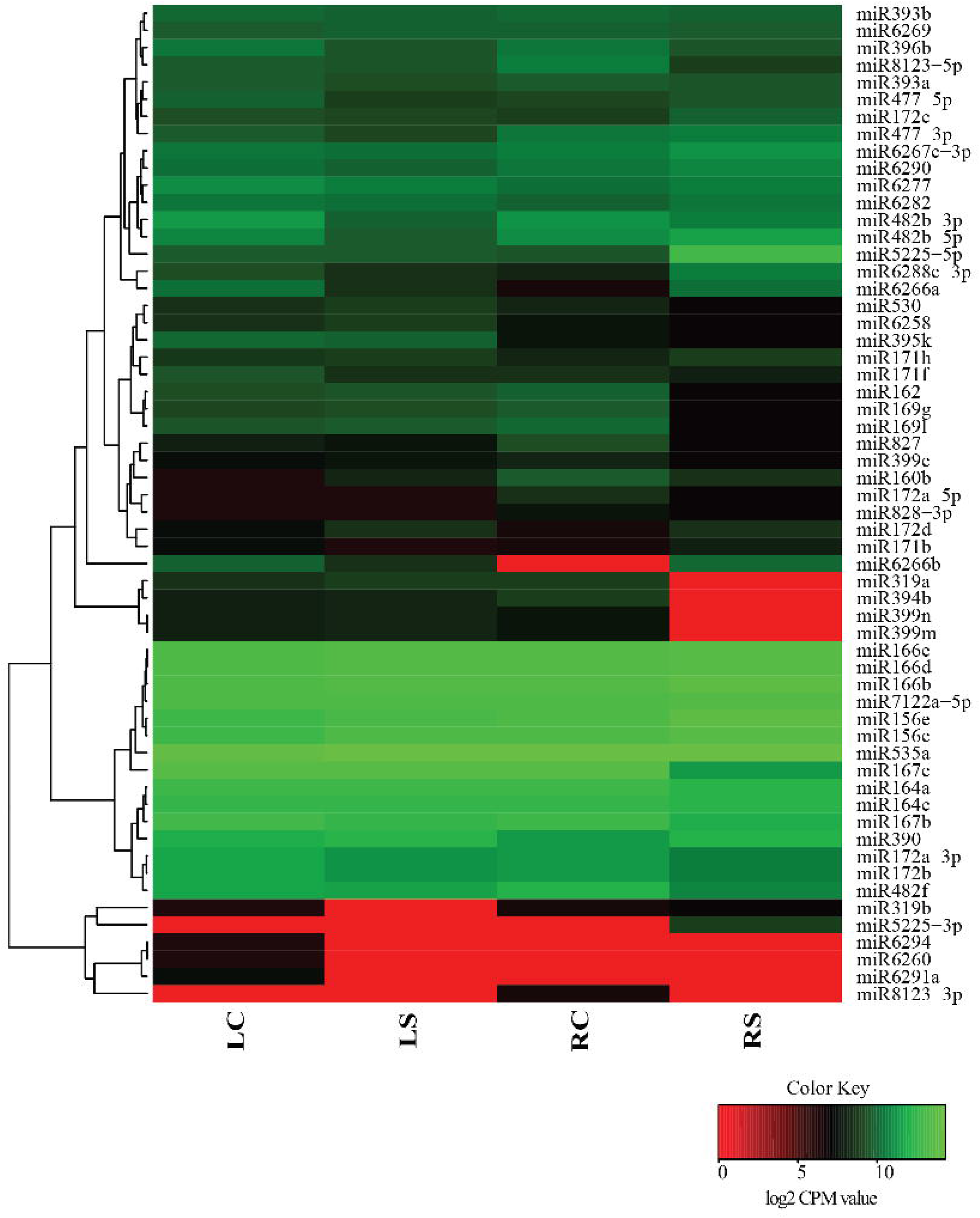
Heat map of TE-related miRNAs. Drought-responsive TE-miRNAs, which were differentially expressed in root and leaf upon, stress treatment represented.

**Table 1.** Drought responsive TE-miRNAs and their fold changes in four sRNA libraries. FC; Fold Change, LC; Leaf-control, LS; Leaf-stress, RC; Root-control, RS; Root-stress. miR6284, which is written in red, is TE-derived, others are TE-related.

| miRNA | FC<br>LC vs LS | FC<br>RC vs RS | Related transposable element |
| --- | --- | --- | --- |
| miR160b | 1,65 | -1,22 | (TGGA)n, LTR_Gm_17_5361, (CATATA)n,<br>Bd1_RLC_72 |
| miR162 | 0,25 | -2,96 | LTR_Sb_chr_02_1099 |
| miR167c | 0,07 | -2,35 | LTR_AC195581.3_5451, LTR_SI_chr_11_2076,<br>LTR_Gm_14_4344, trep4247 |
| miR169g | 0,28 | -2,80 | LTR_AC187047.4_14371 |
| miR169l | 0,23 | -3,22 | (GA)n |
| miR172a-5p | 0,06 | -1,63 | trep4247, trep4410, LTR_AC185219.3_1390,<br>trep4410 |
| miR172d | 1,06 | 1,95 | RLC_Barbara_3B_045_E05-1,<br>LTR_Sb1_00020_929, trep4411 |
| miR5225-5p | 0,06 | 3,45 | LTR_SI_chr_00_49 |
| miR6266a | -1,75 | 3,69 | DTC_unnamedfam16_3B_079_D10-1,<br>LTR_Gm_07_2481, (TG)n, trep3525 |
| miR6284 | -2,52 | 2,37 | LTR_SI_chr_06_1092 |
| miR6288c-3p | -0,75 | 2,48 | LTR_SI_chr_05_1081 |
| miR8123-5p | -0,26 | -1,63 | LTR_Gm_09_3151 |
| miR827 | -0,52 | -2,44 | LTR_Sb_chr_06_655, Gypsy-3_Pru-LTR |

A total of 346 target transcripts regulated by TE-miRNAs were predicted by psRNAtarget tool (Supplementary Table 2). These were subjected to functional annotation with B2GO resulting a total of 564 GO terms (Supplementary Table 3). Biological process of target transcripts, regulated by TE-miRNAs, is mainly regulation of transcription (Supplementary Table 3). Fifty-five of total 355 biological process hits are relevant to regulation of transcription. Oxidation-reduction process, auxin signaling pathway, phosphorylation, translation and response to stimuli are the other mostly represented processes. Response to stimuli includes; abscisic acid, auxin, oxidative stress, salt stress, water deprivation, osmotic stress responses. In the case of molecular function, the targets generally role in binding, mostly DNA and RNA binding, 62 hits were found as either DNA or RNA binding. Hydrolase and transferase activities, ion binding and kinase activity are also prevalent in total 418 molecular function hits. Membrane, chloroplast and nucleus are the most common locations of the targets. Among all targets, maximum number of GO terms is hit to the sequence DW345250, histidine kinase 1, a target of miR482-5p (Supplementary Table 2). Some of the hits are related with biological process, response to osmotic stress, water deprivation, and seed maturation. As molecular function, the hits are, osmosensor and kinase activity. The target transcripts of drought responsive TE-miRNAs are related with transcription, translation, plant development, auxin and abscisic acid signaling pathways and cell division mostly, in terms of biological process (Supplementary Table 3).

## DISCUSSION

In this study we revealed a total 63 TE-miRNAs (either TE-derived or TE-related) in *P. persica* genome, and analyzed their tissue-specific expression pattern in response to drought. We identified *PpeMIR6284* as a TE-derived miRNA gene since a considerable part of (74%) gene sequence is located in the long terminal repeat of a retrotransposon. This finding suggests that, while this retrotransposon translocates by a copy-paste mechanism, it might possibly carry the precursor of miR6284 within the transposon itself.

LTRs are one of the most common types of TEs, which are potentially located in the 1 kb region of TE-related miRNAs. The LTRs or any other TEs co-located with miRNA genes in a 1 kb region, might aid these miRNA genes to move while jumping from one location to the other in the genome. Some members of miR166, miR171, miR172, miR399, miR482 families were detected in the 1 kb region of TEs, thus it might denote that, these families have been affected from translocating events of TEs nearby.

The expression of TE-derived miR6284 in leaves was detected as suppressed 2.5 fold whereas it was upregulated nearly 2.4 fold in root upon drought (Figure 3). Tissue-specific differential regulation of the TE-derived miR6284 was observed in response to drought. Additionally, we discovered that, the expression level of many of the drought responsive TE-miRNAs (FC>1.5) were differentially regulated only in roots. For miR169l, as an example, the root fold change is -3.2 but there is no significant change in the leaf tissue. Only two TE-miRNAs (miR6266a and miR160b) were regulated in leaf with fold changes of -1.7 and 1.6 respectively. On the other hand, 7 of the predicted TE-related miRNA precursor transcripts (MIR399a, MIR6267c, MIR6288c, MIR6291b, MIR6296, MIR7122a and MIR828) were found to be undifferentiated upon drought treatment.

Blast2GO results indicated that most of the TE-miRNA target transcripts are functionally related with transcription mechanism in terms of biological process, and DNA or RNA and binding as molecular function (Supplementary Table 3). Therefore it can be deduced that; TE-related miRNAs target generally transcription factors. Auxin related processes are also common among the GO terms of target transcripts. Since, it is a plant growth hormone involved in developmental, it can also regulate transcription (.Perrot-Rechenmann 2014). Some of the target transcripts found to be related to developmental processes and cell growth independent of auxin implying that, TE-related miRNAs in peach might target the mRNAs of the genes to regulate the growth and development. TE-miRNAs might also control the process of response to various stimuli including osmotic stress, water deprivation and oxidative stress by targeting the transcripts associate with these mechanisms. Expression analysis and GO outputs indicated that drought responsive TE-related miRNAs might regulate stress tolerance by targeting the genes encoding transcription factors and proteins to control plant development either auxin dependent or independent in tissue specific manner.

## Supplementary materials

Supplementary table 1.TE-related miRNAs & TEs in 1kb window

Supplementary table 2.TE-miRNA targets

Supplementary table 3. Blast2GO table

Supplementary table 4. Blast2GO table of the target transcripts of drought responsive TE-miRNAs

## REFERENCES

Abrouk M, Zhang R, Murat F, Li A, Pont C, Mao L, Salse J (2012) Grass microRNA gene paleohistory unveils new insights into gene dosage balance in subgenome partitioning after whole-genome duplication. The Plant Cell 24:1776–1792

Akdogan G, Tufekci ED, Uranbey S, Unver T (2016) miRNA-based drought regulation in wheat. Funct Integr Genomics 16:221–233. DOI 10.1007/s10142-015-0452-1

Bennetzen JL, Wang H (2014) The contributions of transposable elements to the structure, function, and evolution of plant genomes. Annual review of plant biology 65:505–530. DOI 10.1146/annurev-arplant-050213-035811

Conesa A, Götz S, García-Gómez JM, Terol J, Talón M, Robles M (2005) Blast2GO: a universal tool for annotation, visualization and analysis in functional genomics research. Bioinformatics 21:3674–3676

Dai X, Zhao PX (2011) psRNATarget: a plant small RNA target analysis server. Nucleic acids research 39:W155–W159

Deragon J, Casacuberta J, Panaud O (2008) Plant transposable elementsPlant Genomes. Karger Publishers, pp. 69–82.

Eldem V, Celikkol Akcay U, Ozhuner E, Bakir Y, Uranbey S, Unver T (2012) Genome-wide identification of miRNAs responsive to drought in peach (Prunus persica) by high-throughput deep sequencing. PLoS One 7:e50298. DOI 10.1371/journal.pone.0050298

Eldem V, Okay S, ÜNVER T (2013) Plant microRNAs: new players in functional genomics. Turkish Journal of Agriculture and Forestry 37:1–21

Frazier TP, Sun G, Burklew CE, Zhang B (2011) Salt and drought stresses induce the aberrant expression of microRNA genes in tobacco. Molecular biotechnology 49:159–165

Friedländer MR, Mackowiak SD, Li N, Chen W, Rajewsky N (2012) miRDeep2 accurately identifies known and hundreds of novel microRNA genes in seven animal clades. Nucleic acids research 40:37–52

Hajyzadeh M, Turktas M, Khawar KM, Unver T (2015) miR408 overexpression causes increased drought tolerance in chickpea. Gene 555:186–193. DOI 10.1016/j.gene.2014.11.002

Kantar M, Unver T, Budak H (2010) Regulation of barley miRNAs upon dehydration stress correlated with target gene expression. Funct Integr Genomics 10:493–507. DOI 10.1007/s10142-010-0181-4

Kejnovsky E, Hawkins JS, Feschotte C (2012) Plant transposable elements: biology and evolutionPlant Genome Diversity Volume 1. Springer, pp. 17–34.

Kersey PJ, Allen JE, Armean I, Boddu S, Bolt BJ, Carvalho-Silva D, Christensen M, Davis P, Falin LJ, Grabmueller C (2016) Ensembl Genomes 2016: more genomes, more complexity. Nucleic acids research 44:D574–D580

Kozomara A, Griffiths-Jones S (2014) miRBase: annotating high confidence microRNAs using deep sequencing data. Nucleic acids research 42:D68–D73

Li Y, Li C, Xia J, Jin Y (2011) Domestication of transposable elements into MicroRNA genes in plants. PloS one 6:e19212. DOI 10.1371/journal.pone.0019212

Lisch D (2013) How important are transposons for plant evolution? Nature reviews Genetics 14:49–61. DOI 10.1038/nrg3374

Maher C, Stein L, Ware D (2006) Evolution of Arabidopsis microRNA families through duplication events. Genome research 16:510–519. DOI 10.1101/gr.4680506

Mallory AC, Bartel DP, Bartel B (2005) MicroRNA-directed regulation of Arabidopsis AUXIN RESPONSE FACTOR17 is essential for proper development and modulates expression of early auxin response genes. The Plant Cell 17:1360–1375

Mallory AC, Dugas DV, Bartel DP, Bartel B (2004) MicroRNA regulation of NAC-domain targets is required for proper formation and separation of adjacent embryonic, vegetative, and floral organs. Current Biology 14:1035–1046

Palatnik JF, Allen E, Wu X, Schommer C, Schwab R, Carrington JC, Weigel D (2003) Control of leaf morphogenesis by microRNAs. Nature 425:257–263

Perrot-Rechenmann C (2014) Auxin Signaling in PlantsMolecular Biology. Springer, pp. 245–268.

Piriyapongsa J, Jordan IK (2008) Dual coding of siRNAs and miRNAs by plant transposable elements. RNA 14:814–821. DOI 10.1261/rna.916708

Qin F, Shinozaki K, Yamaguchi-Shinozaki K (2011) Achievements and challenges in understanding plant abiotic stress responses and tolerance. Plant and Cell Physiology 52:1569–1582

Quinlan AR (2014) BEDTools: the Swiss-army tool for genome feature analysis. Current protocols in bioinformatics:11.12.11-11.12. 34

Santiago N, Herráiz C, Goñi JR, Messeguer X, Casacuberta JM (2002) Genome-wide analysis of the Emigrant family of MITEs of Arabidopsis thaliana. Molecular biology and evolution 19:2285–2293

Sun J, Zhou M, Mao Z, Li C (2012) Characterization and evolution of microRNA genes derived from repetitive elements and duplication events in plants. PloS one 7:e34092. DOI 10.1371/journal.pone.0034092

Wicker T, Sabot F, Hua-Van A, Bennetzen JL, Capy P, Chalhoub B, Flavell A, Leroy P, Morgante M, Panaud O(2007) A unified classification system for eukaryotic transposable elements. Nature Reviews Genetics 8:973–982

Xie F, Stewart CN, Taki FA, He Q, Liu H, Zhang B (2014) High-throughput deep sequencing shows that microRNAs play important roles in switchgrass responses to drought and salinity stress. Plant Biotechnology Journal 12:354–366

Xie F, Wang Q, Sun R, Zhang B (2015) Deep sequencing reveals important roles of microRNAs in response to drought and salinity stress in cotton. Journal of experimental botany 66:789–804

Yan Y, Zhang Y, Yang K, Sun Z, Fu Y, Chen X, Fang R (2011) Small RNAs from MITE-derived stem-loop precursors regulate abscisic acid signaling and abiotic stress responses in rice. The Plant journal: for cell and molecular biology 65:820–828. DOI 10.1111/j.1365-313X.2010.04467.x

Zhang B (2015) MicroRNA: a new target for improving plant tolerance to abiotic stress. Journal of experimental botany 66:1749–1761. DOI 10.1093/jxb/erv013

Zhang X, Zou Z, Gong P, Zhang J, Ziaf K, Li H, Xiao F, Ye Z (2011) Over-expression of microRNA169 confers enhanced drought tolerance to tomato. Biotechnology letters 33:403–409

Zhao B, Liang R, Ge L, Li W, Xiao H, Lin H, Ruan K, Jin Y (2007) Identification of drought-induced microRNAs in rice. Biochemical and biophysical research communications 354:585–590

Zhou M, Li D, Li Z, Hu Q, Yang C, Zhu L, Luo H (2013) Constitutive expression of a miR319 gene alters plant development and enhances salt and drought tolerance in transgenic creeping bentgrass. Plant physiology 161:1375–1391

